# Heterozygous ATP-binding Cassette Transporter G5 Gene Deficiency and Risk of Coronary Artery Disease

**DOI:** 10.1101/780734

**Authors:** Akihiro Nomura, Connor A. Emdin, Hong Hee Won, Gina M. Peloso, Pradeep Natarajan, Diego Ardissino, John Danesh, Heribert Schunkert, Adolfo Correa, Matthew J. Bown, Nilesh J. Samani, Jeanette Erdmann, Ruth McPherson, Hugh Watkins, Danish Saleheen, Roberto Elosua, Masa-aki Kawashiri, Hayato Tada, Namrata Gupta, Svati H. Shah, Daniel J. Rader, Stacey Gabriel, Amit V. Khera, Sekar Kathiresan

## Abstract

**Background:** Familial sitosterolemia is a rare, recessive Mendelian disorder characterized by hyperabsorption and decreased biliary excretion of dietary sterols. Affected individuals typically have complete genetic deficiency – homozygous loss-of-function (LoF) variants — in the ATP-binding cassette transporter G5 (*ABCG5*) or G8 (*ABCG8*) genes, and have substantially elevated plasma sitosterol and low-density lipoprotein cholesterol (LDL-C) levels. The impact of partial genetic deficiency of *ABCG5* or *ABCG8*, as occurs in heterozygous carriers of LoF variants, on LDL-C and risk of coronary artery disease (CAD) has remained uncertain.

**Methods:** We first recruited nine sitosterolemia families, identified causative LoF variants in *ABCG5* or *ABCG8*, and evaluated the associations of these *ABCG5* or *ABCG8* LoF variants with plasma phytosterols and lipid levels. We next assessed for LoF variants in *ABCG5* or *ABCG8* in CAD cases (n=29,361) versus controls (n=357,326). We tested the association of rare LoF variants in *ABCG5* or *ABCG8* with blood lipids and risk for CAD. Rare LoF variants were defined as protein-truncating variants with minor allele frequency less than 0.1% in *ABCG5* or *ABCG8*.

**Results:** In sitosterolemia families, seven pedigrees harbored causative LoF variants in *ABCG5* and two pedigrees in *ABCG8*. Homozygous LoF variants in either *ABCG5* or *ABCG8* led to marked elevations in sitosterol and LDL-C. Of those sitosterolemia families, heterozygous carriers of *ABCG5* LoF variants exhibited increased sitosterol and LDL-C levels compared to non-carriers. Within the large-scale CAD case-control cohorts, prevalence of rare LoF variants in *ABCG5* and in *ABCG8* were approximately 0.1% each. *ABCG5* heterozygous LoF variant carriers had significantly elevated LDL-C levels (24.7 mg/dL; 95% confidence interval [CI] 14 to 35; P=1.1×10^−6^), and were at two-fold increased risk of CAD (odds ratio 2.06, 95% CI 1.27 to 3.35; P=0.004). By contrast, *ABCG8* heterozygous LoF carrier status was not associated with increased LDL-C or risk of CAD.

**Conclusions:** Although familial sitosterolemia is traditionally considered as a recessive disorder, we observed that heterozygous carriers of a LoF variant in *ABCG5* had significantly increased sitosterol and LDL-C levels and a two-fold increase in risk of CAD.

## Introduction

Familial sitosterolemia (OMIM #210250) is a rare Mendelian recessive disorder characterized by tendonous xanthomas^1^, high plasma plant sterols and cholesterol levels, and increased risk of premature myocardial infarction.^2–4^ The ATP-binding cassette transporters G5 (*ABCG5*) and G8 (*ABCG8*) are primary causal genes of familial sitosterolemia. *ABCG5, ABCG8*, and N-terminal Niemann-Pick C1 Like 1 (*NPC1L1*)^5, 6^ determine the efflux and absorption of sterols on the surface of intestine and bile duct.^7, 8^ *NPC1L1* regulates sterol absorption whereas *ABCG5* and *ABCG8* form obligate heterodimers^9^ and coordinately control the excretion at both the brush border membrane of enterocyte and the apical membrane of hepatocytes.^5, 10-12^

Complete deficiency due to homozygous or compound heterozygous loss-of-function (LoF) variants in *ABCG5* and/or *ABCG8* causes markedly increased sitosterolemia and cholesterol levels, and potentially accelerated atherosclerotic disease as well.^1–4^ Genome-wide association studies also demonstrated that loci at the *ABCG5*-*ABCG8* gene region were associated with phytosterols, low-density lipoprotein (LDL) cholesterol,^13^ and risk of coronary artery disease (CAD).^14^ However, it is uncertain whether partial deficiency of *ABCG5 or ABCG8* as conferred by LoF variants are also associated with higher cholesterol levels and an increased risk of CAD.

Here, we explored the metabolic consequences of *ABCG5* or *ABCG8* deficiencies. We recruited probands and relatives in sitosterolemia families and assessed whether observed *ABCG5* or *ABCG8* causative LoF variants were associated with increased plasma phytosterols and LDL cholesterol. We then analyzed exome sequences from 112,700 participants and genotype data from an additional 293,134 individuals to test whether carriers of rare heterozygous LoF variants in *ABCG5* or *ABCG8* had elevated blood lipids and risk of CAD.

## Methods

### Study Participants

Nine sitosterolemia pedigrees with phytosterol and lipid profiles derived from the Kanazawa University Mendelian Disease (KUMD) Registry were evaluated for *ABCG5* and *ABCG8* variants (**Supplemental Figure 1**).^3^ Sitosterolemia was diagnosed by 1) plasma sitosterol concentration ≥10 μg/mL and 2) presence of tendon or tuberous xanthomas and/or history of premature CAD.^15^ Causative LoF variants in the pedigrees were defined as pathogenic or likely pathogenic by the American College of Medical Genetics (ACMG) standard criteria,^16^ and were identified by Sanger sequencing or whole exome sequencing (WES). Controls were unaffected relatives without any causative LoF variants in the sitosterolemia pedigrees.

Next, *ABCG5* and *ABCG8* were sequenced in the Myocardial Infarction Genetics consortium (MIGen), UK Biobank and the TruSeq Custom Amplicon target resequencing (TSCA) studies. MIGen case control studies included the Italian Atherosclerosis Thrombosis and Vascular Biology (ATVB) study,^17^ the Deutsches Herzzentrum München Myocardial Infarction Study (DHM),^18^ the Exome Sequencing Project Early-Onset Myocardial Infarction study (ESP-EOMI),^19^ the Jackson Heart Study (JHS),^20^ the Leicester Acute Myocardial Infarction Study (Leicester),^21^ the Lubeck Myocardial Infarction Study (Lubeck),^22^ the Ottawa Heart Study (OHS),^23^ the Precocious Coronary Artery Disease (PROCARDIS) study,^24^ the Pakistan Risk of Myocardial Infarction Study (PROMIS),^25^ and the Registre Gironi del COR (Gerona Heart Registry or REGICOR) study.^26^ TSCA included the Duke CATHGEN study (DUKE),^27^ the MedStar study (MedStar),^28^ and the PennCath study (PennCath).^28^ An additional 293,134 individuals in UK Biobank underwent array based genotyping for the *ABCG5* stop variant rs199689137 (p.R446Ter) and were included in the analysis.

All participants in each study provided written informed consent for genetic studies. The institutional review board at Partners HealthCare (Boston, MA, USA) and each participating institution approved the study protocol.

### Lipid measurement and coronary artery disease ascertainment

In sitosterolemia pedigree-based analysis, all blood samples were obtained after a 12-hour overnight fast, either before initiation of lipid-lowering treatment or after discontinuation of medication for at least 4 weeks. Plasma levels of non-cholesterol sterols were determined using gas– liquid chromatography–mass spectrometry.^4^ In addition to absolute non-cholesterol sterol levels, cholesterol adjusted ratios (each non-cholesterol sterol level to total cholesterol [TC] level ratio) were also evaluated. Plasma concentrations of TC, triglyceride, and HDL cholesterol were determined in MIGen, TSCA and UK Biobank using enzymatic assays. In MIGen and TSCA, LDL cholesterol level was calculated using the Friedewald equation for those with triglycerides <400 mg/dL. In the UK Biobank, LDL cholesterol levels were directly measured using an antibody-based assay.

In MIGen, CAD cases were identified as early-onset (premature) myocardial infarction (defined as ≤ 50 years old in male and ≤ 60 years old in female). In the UK Biobank, CAD cases were defined as myocardial infarction at any age. In TSCA, coronary artery disease was defined as stenosis on angiography (≥ 1 coronary vessel with > 50% stenosis) at ages ≤ 50 years old for men and ≤ 60 years old for women.

### Exome sequencing

WES or targeted sequencing for KUMD, MIGen, and TSCA was performed at the Broad Institute as previously described.^5,6^ In brief, genomic deoxyribonucleic acid was captured on protein-coding regions using the NimbleGen Sequencing Capture Array or the Illumina TrueSeq Custom Amplicon. Sequencing reads were aligned to a human reference genome (build 37) using the Burrows–Wheeler Aligner-Maximal Exact Match algorithm. Aligned non-duplicate reads were locally realigned, and base qualities were recalibrated using the Genome Analysis ToolKit (GATK) software.^29^ Variants were jointly called using the GATK HaplotypeCaller software. WES in UK Biobank was performed by the Regeneron Genetics Center as previously described.^7^

In large cohort studies, rare LoF variants were defined as those with minor allele frequency less than 0.1% and which caused: (1) insertions or deletions of DNA that modified the reading frame of protein translation (frameshift); (2) point mutations at conserved splice site regions that altered the splicing process (splice-site); or (3) point mutations that changed an amino acid codon to a stop codon, leading to the truncation of a protein (nonsense). LoF variants were identified using the LOFTEE plugin of the Variant Effect Predictor software (version 82). ^8, 9^

### Statistical Analysis

In sitosterolemia family-based analyses, the differences in lipid levels, lipoproteins and non-cholesterol sterols by *ABCG5* or *ABCG8* LoF variant carrier status were assessed using the *Mann-Whitney U* test. The effects of *ABCG5* and *ABCG8* rare LoF variants on lipid profiles in MIGen, UK Biobank and TSCA was evaluated using linear regression, adjusting for age, gender, study, and first five principal components of ancestry. A Cochran–Mantel–Haenszel statistic meta-analysis for stratified 2-by-2 tables was used to associate *ABCG5* and *ABCG8* rare LoF variants with risk of CAD. As a sensitivity analysis, we performed an inverse variance weighted fixed effects meta-analysis of the adjusted odds ratio, derived in each cohort using logistic regression adjusted for age, sex, study and five principal components of ancestry. P values of less than 0.025 were considered to indicate statistical significance (i.e., Bonferroni correction for the testing of two genes). Statistical analyses were performed using R software version 3.4.3 (The R Project for Statistical Computing, Vienna, Austria).

## Results

### *ABCG5* or *ABCG8* causative LoF variants, blood phytosterol and cholesterol levels in sitosterolemia families

We recruited nine Japanese families with sitosterolemia and sequenced the exons of the *ABCG5* and *ABCG8* genes in 47 individuals from these families (**Supplemental Figure 1**). Among the individuals within these families, 9 carried a homozygous or compound heterozygous *ABCG5* or ABC*G8* causative LoF variants while 28 carried a heterozygous *ABCG5* or ABC*G8* LoF causative variants. Most of those LoF variants were considered as pathogenic protein truncating or missense variants by the ACMG standard criteria (**Supplemental Table 1**). As expected, *ABCG5* and *ABCG8* homozygote or compound heterozygous LoF variant carriers showed very high sitosterol / TC ratio and LDL cholesterol levels compared to non-carriers. Regarding heterozygous state, carriers of *ABCG5* or *ABCG8* heterozygous LoF variant carriers exhibited increased sitosterol / TC ratio compared with non-carriers. Moreover, *ABCG5* heterozygous LoF variant carrier status was associated with an increased LDL cholesterol level. (Table 1 and Figure 1).

**Figure 1.**
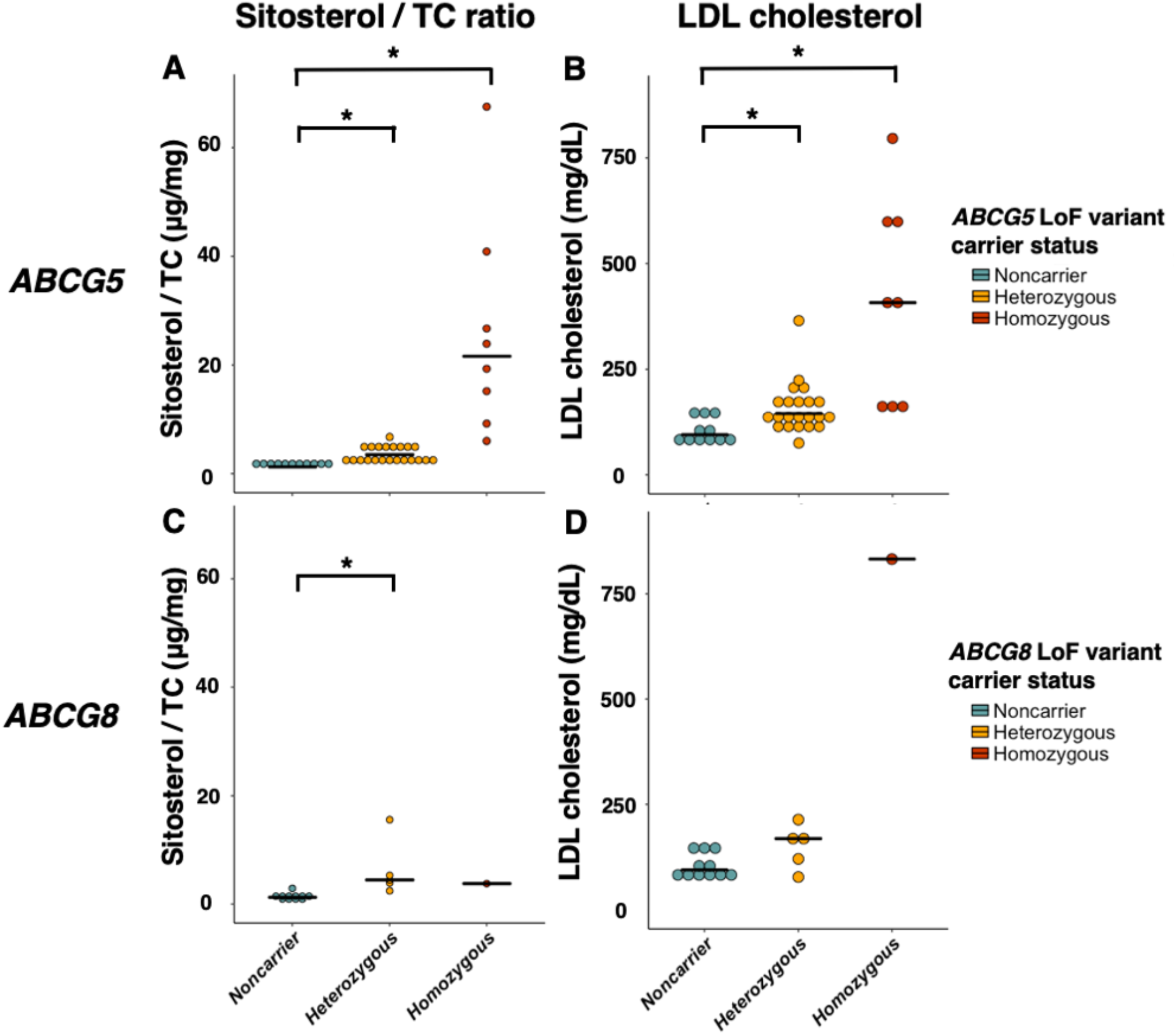
Sitosterol to total cholesterol ratio and LDL cholesterol levels among individuals with homozygous and heterozygous sitosterolemia, and unaffected controls in sitosterolemia families. Each dot indicates an individual’s value. Each horizontal line represents a mean value for each carrier status. *: P < 0.025 by *Mann-Whitney U* test.

**Table 1.** Clinical characteristics by *ABCG5* and *ABCG8* variant carrier status in sitosterolemia families.

|  | Non-carrier | *ABCG5* |  | *ABCG8* |  |
| --- | --- | --- | --- | --- | --- |
|  |  | Hetero | Homo | Hetero | Homo |
| N | 11 | 22 | 8 | 5 | 1 |
| Age, mean ± SD | 40.5 ± 19 | 41.1 ± 21 | 12.9 ± 20 | 45.2 ± 19 | 1 |
| Male sex, n (%) | 3 (27) | 12 (55) | 3 (38) | 2 (40) | 0 |
| Lipid profile |  |  |  |  |  |
| Total cholesterol, mg/dL, mean ± SD | 180 ± 28 | 236 ± 70* | 512 ± 250* | 258 ± 67 | 968 |
| LDL cholesterol, mg/dL, mean ± SD | 105 ± 30 | 158 ± 59* | 412 ± 240* | 150 ± 52 | 832 |
| HDL cholesterol, mg/dL, mean ± SD | 61.1 ± 15 | 56.1 ± 19 | 47.4 ± 13 | 57.0 ± 13 | 46 |
| Triglyceride, mg/dL, median (IQR) | 87 (74–96) | 76 (55–150) | 188 (140–248)* | 154 (73–154) | 71 |
| Lipoproteins |  |  |  |  |  |
| Apolipoprotein A1, mg/dL, median (IQR) | 150 (137–165) | 140 (128–150) | 106 (97–129) | NA | NA |
| Apolipoprotein B, mg/dL, median (IQR) | 67 (63–93) | 106 (91–117) | 262 (198–303)* | NA | NA |
| Non-cholesterol sterols |  |  |  |  |  |
| Sitosterol, μg/mL, median (IQR) | 2.2 (1.8–2.7) | 7.6 (5.5–10)* | 102 (74–125)* | 9.9 (8.2–12)* | 36.5 |
| Campesterol, μg/mL, median (IQR) | 3.6 (3.2–4.5) | 13 (10–14)* | 70 (65–95)* | NA | NA |
| Sitosterol / TC, μg/mg, median (IQR) | 1.2 (1.1–1.4) | 3.5 (2.5–4.6)* | 22 (14–30)* | 4.4 (3.9–5.3)* | 3.8 |
| Campesterol / TC, μg/mg, median (IQR) | 2.2 (1.7–2.6) | 5.2 (4.6–6.5)* | 13 (7.5–18)* | NA | NA |

\*: P value < 0.025 compared to non-carrier controls. P values were calculated by *Mann-Whitney U* test. Abbreviations: IQR, interquartile range; SD, standard deviation; TC, total cholesterol.

### *ABCG5* or *ABCG8* rare heterozygous LoF variation, blood lipids and risk for CAD in large cohorts

Next, we examined whether rare heterozygous LoF variant carrier status in *ABCG5* or *ABCG8* increased blood lipids and were at elevated risk of CAD using large-scale genetics cohorts. We sequenced the protein coding regions of *ABCG5* and *ABCG8* in 112,700 individuals from three datasets: 58,791 participants from MIGen, 52,195 participants in UK Biobank and 1,714 participants from TSCA (Table 2). We detected 108 individuals harboring rare *ABCG5* LoF alleles and the prevalence of *ABCG5* heterozygous carrier status was 0.1%. (**Supplemental Table 2**). We also discovered 154 individuals who harbored rare *ABCG8* LoF alleles, a heterozygous carrier prevalence also around 0.1%.

**Table 2.** Clinical characteristics of participants in MIGen, UK Biobank and targeted resequencing study.

|  | **Myocardial**<br>**Infarction**<br>**Genetics**<br>**Consortium** | **UK Biobank** | **Target**<br>**Resequencing**<br>**Study** |
| --- | --- | --- | --- |
|  | **N = 58,791** | **N = 336,391** | **N = 1,714** |
| Age, years (SD) | 54 (10) | 57 (8) | 57.2 (13) |
| Male gender, n (%) | 41,203 (71) | 156,112 (46) | 1,140 (67) |
| BMI (SD), kg/m<sup>2</sup> | 27 (5) | 27 (5) | 29 (6) |
| Current smoker, n (%) | 18,173 (33) | 25,802 (8) | 639 (37) |
| **Medical history** |  |  |  |
| Coronary artery disease, n (%) | 16,106 (33) | 12,073 (3) | 1,184 (69) |
| Hypertension, n (%) | 16,839 (35) | 153,535 (46) | 798 (47) |
| Type 2 diabetes, n (%) | 11,245 (22) | 15,770 (5) | 277 (16) |
| **Lipid profile** |  |  |  |
| Total cholesterol, mg/dl (SD) | 178 (60) | 222 (41) | 182 (51) |
| LDL cholesterol, mg/dl<sup>†</sup> (SD) | 109 (45) | 138 (34) | 108 (41) |
| HDL cholesterol, mg/dl (SD) | 36 (14) | 56 (15) | 43 (7) |
| Triglycerides, mg/dl (SD) | 154 (127) | 155 (90) | 135 (82) |

Abbreviations: SD, standard deviation.

Individuals carrying *ABCG5* LoF variants had significantly increased total cholesterol levels (17 mg/dl; 95% confidence interval [CI], 13 to 32; P = 6.9×10^−6^) and LDL cholesterol levels (25 mg/dl, 95% CI, 13 to 32; P = 1.1×10^−6^, Figure 2). Then, we investigated the association between rare *ABCG5* heterozygous LoF variant carrier status and CAD risk using more than 380,000 participants from the three WES cohorts and additional UK biobank genotyping array-based cohort. We identified 34 carriers of *ABCG5* heterozygous LoF variants among 29,321 coronary artery disease cases (0.1%) and 63 among 357,326 controls (0.02%). In a Cochran-Mantel-Haenszel fixed-effect meta-analysis, individuals carrying *ABCG5* heterozygous LoF variants were at two-fold risk of CAD (Odds ratio [OR], 2.06; 95% CI, 1.27 to 3.35; P value = 0.004) (Figure 3). A similar effect estimate was noted in a meta-analysis of adjusted odds ratios derived using logistic regression (OR 2.04; 95% CI 1.28 to 3.26; P = 0.003).

**Figure 2.**
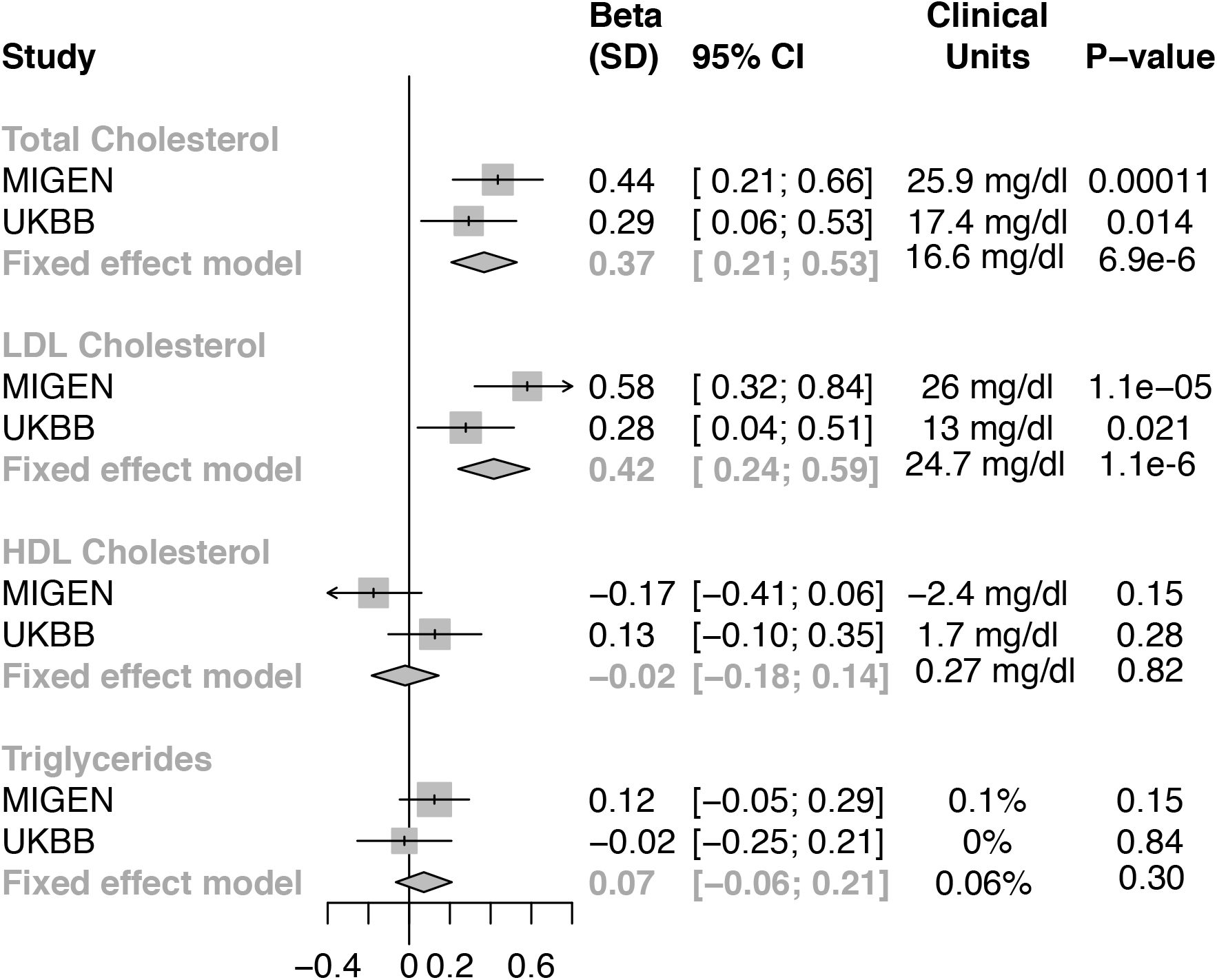
Effects of loss-of-function variants in *ABCG5* on blood lipid profiles from MIGen and UK Biobank. Effect sizes were calculated using linear regression adjusted by age, gender, study, case-control status, and first five principal components of ancestry. Triglyceride was natural log-transformed before analysis. Fixed-effects meta-analysis was applied to combine results.

**Figure 3.**
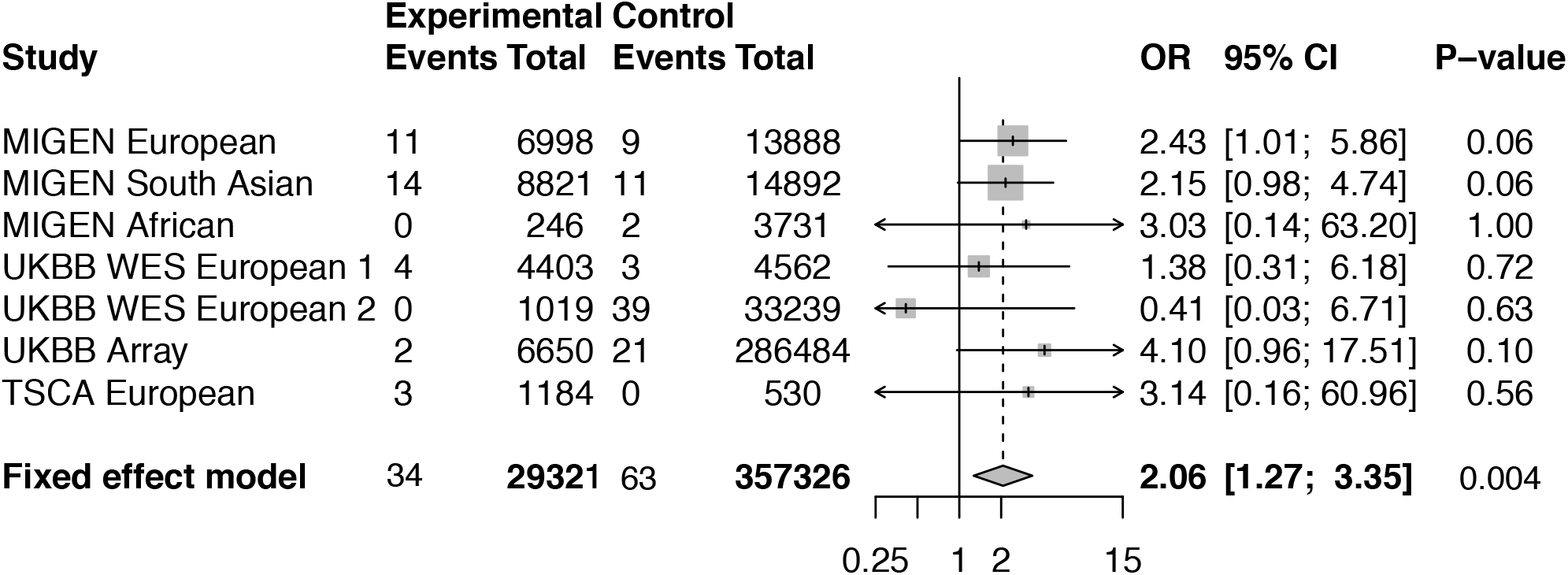
Effect of loss-of-function variants in *ABCG5* on myocardial infarction risk. A meta-analysis across studies was performed using the Cochran–Mantel–Haenszel statistics for stratified 2-by-2 tables. MIGen, myocardial infarction genetics consortium.

In contrast to *ABCG5*, carriers of rare *ABCG8* heterozygous LoF variants did not exhibit significant increase in any of blood lipids including LDL cholesterol (beta, 0.08; 95% CI, −0.06 to 0.23; P = 0.25) (**Supplemental Table 3**). Moreover, *ABCG8* heterozygous LoF variant carrier status was not at elevated risk for CAD (OR, 0.79; 95% CI, 0.47 to 1.35; P = 0.39) (**Supplemental Table 3**).

We explored whether the effect size of *ABCG5* LoF variants on CAD risk was consistent with their effect on LDL cholesterol. We observed a linear dose-response relationship between CAD risk and LDL cholesterol change conferred by DNA sequence variants in *LDLR, PCSK9*, *ABCG5* and *ABCG8*. The effect of *ABCG5* LoF variants on CAD risk (106% increase in risk) was consistent with the estimate based on the change in LDL cholesterol (25 mg/dl) (Figure 4).

**Figure 4.**
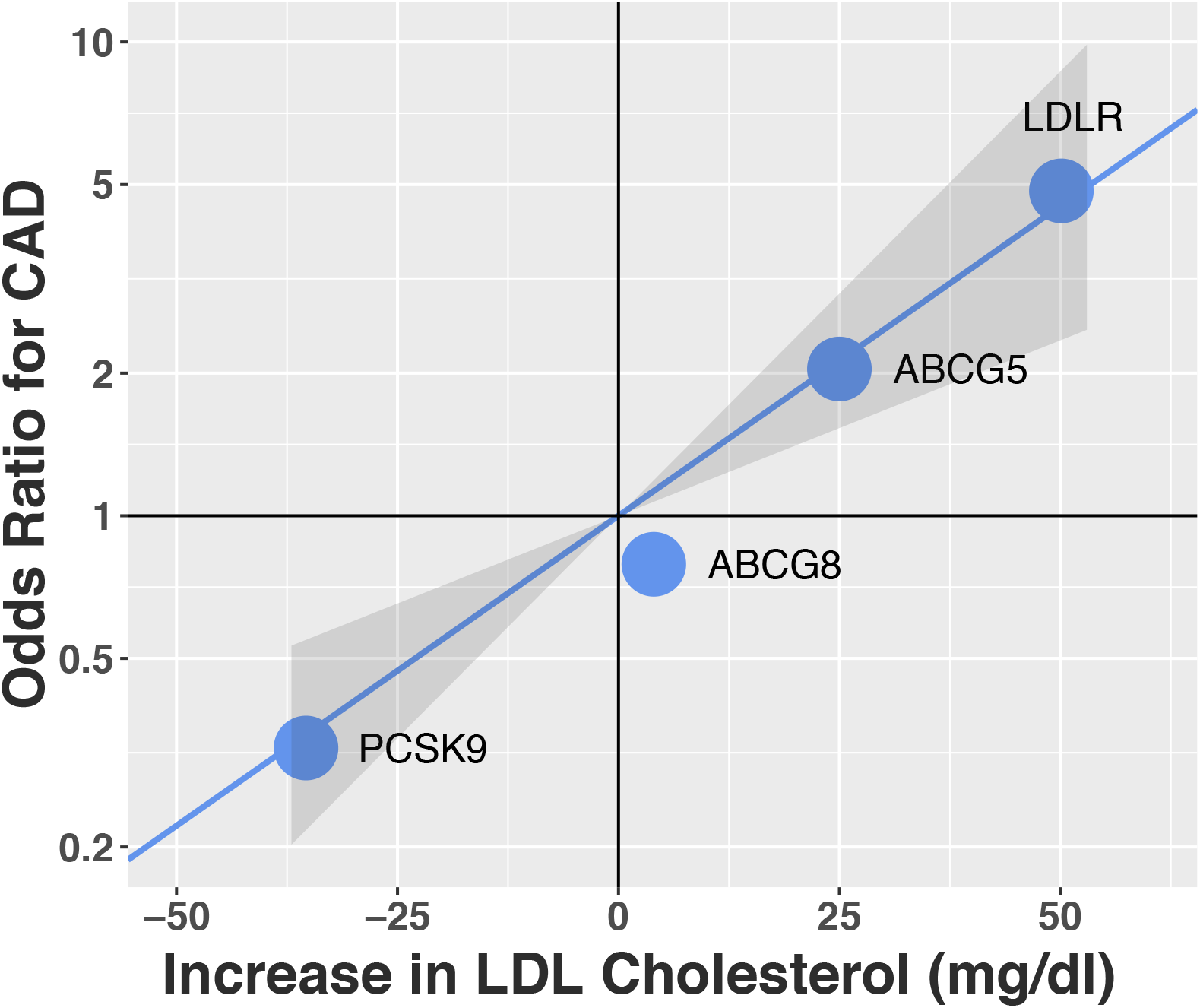
Relationship between effect on LDL cholesterol levels and coronary artery disease risk for *ABCG5*, *ABCG8*, *PCSK9* and LDLR loss-of-function variants. CAD, coronary artery disease; LDL, low-density lipoprotein

## Discussion

In this study, we evaluated whether rare heterozygous LoF variations in *ABCG5* or *ABCG8* were associated with blood lipid levels and CAD risk. We used two different approaches — sitosterolemia family-based analysis and population-based analyses from over 380,000 individuals — to test whether rare heterozygous LoF variants in *ABCG5* or *ABCG8* associated with phytosterols, lipids and CAD. We found that carriers of heterozygous LoF variants in *ABCG5* had higher sitosterol and ~25 mg/dl higher LDL cholesterol and were at two-fold risk of CAD.

These results permit several conclusions. First, individuals who carried rare heterozygous LoF variants in *ABCG5* had significantly elevated LDL cholesterol levels and were at elevated risk for CAD. Although there have been reports of premature atherosclerosis among patients with homozygous sitosterolemia,^10^ it was unclear if *ABCG5* partial deficiency by heterozygous LoF variants also increases blood lipid levels and CAD risk. The findings of this study clearly demonstrated that *ABCG5* partial deficiency increased both LDL cholesterol level and CAD risk. Considering the higher burden of CAD at a population level, these findings suggests that *ABCG5* LoF variant carriers may derive clinical benefit from LDL cholesterol lowering therapy. Importantly, the *NPC1L1* inhibitor ezetimibe is known to reduce intestinal cholesterol and phytosterol absorption in patients with sitosterolemia, and could have increased efficacy in individuals with partial *ABCG5* deficiency.^30^

Second, it has been unclear whether elevated plant sterol levels or elevated blood cholesterol levels cause atherosclerosis among patients with sitosterolemia.^14^ The proportional relationship between CAD risk and elevation in LDL cholesterol levels observed among *ABCG5* heterozygous LoF carriers suggests that the substantial increase in blood LDL cholesterol levels among individuals with sitosterolemia causes atherosclerosis (Figure 3). These findings are also consistent with a recent meta-analysis that did not observe a significant association between circulating sitosterol levels and risk of cardiovascular disease.^31^ Moreover, the effect size of *ABCG5* heterozygous LoF variant carrier status on both blood lipids and CAD risk was consistent with predictions based on known familial hypercholesterolemia and hypobetalipoproteinemia variants (Figure 4). This result is consistent with recent studies, and it might support the hypothesis that the main driver of CHD risk in *ABCG5* LoF variant carriers was LDL cholesterol rather than plant sterols.

This study has several limitations. First, functional analyses of each LoF variant were not performed. Consequently, some annotated LoF variants included in this study may not actually cause deficiency in the *ABCG5*/*G8* proteins to increase plant sterols and blood lipids. This would underestimate of the effect of *ABCG5* and especially *ABCG8* deficiency on blood lipid levels and CAD risk. Second, CAD definition was different among study cohorts. However, the effect direction among studies was largely consistent and we observed little heterogeneity for the meta-analysis (I squared of 0%).

In conclusion, 0.1% of population carried rare LoF variants in *ABCG5* and these heterozygous carriers of had an elevated sitosterol and LDL cholesterol levels and were at two-fold risk for CAD.

## Acknowledgements

We would like to express our gratitude to all the participants and staff of KUMD, MIGen consortium, TSCA, and UK biobank for their outstanding contributions.

