## Supplemental data for "Heterozygous ATP-binding Cassette Transporter G5 Gene Deficiency and Risk of Coronary Artery Disease"

### Table of Contents

**Supplemental Table 1. List of *ABCG5* rare loss-of-function variants from sitosterolemia family analysis.**

**Supplemental Table 2. Rare loss-of-function variants identified in *ABCG5*.**

**Supplemental Table 3. Association of *ABCG8* loss-of-function variants with blood lipids and coronary artery disease risk among three WES cohorts.**

**Supplemental Figure 1. Pedigrees of sitosterolemia families.**

**Supplemental Table 1. List of *ABCG5* and *ABCG8* rare loss-of-function variants from sitosterolemia family analysis.**

| Variant (GRCh37) | Consequence | Family # | ACMG guideline classification |
|---|---|---|---|
| ***ABCG5*** | | | |
| 2: 44040449_C/T | Splice acceptor | 5 | Pathogenic (PVS1+PS3) |
| 2: 44041700_TAAAAG/T | Frameshift: P558fsTer | 1,3 | Pathogenic (PVS1+PS3) |
| 2: 44050063_G/A | Premature stop: R446X | 3 | Pathogenic (PVS1+PS3) |
| 2: 44051120_C/T | Missense: R419H | 5 | Pathogenic (PS1+PS3) |
| 2: 44051210_C/T | Missense: R389H | 1,2,4,6 | Pathogenic (PS1+PS3) |
| 2: 44052028_T/C | Missense: M302V | 2 | Pathogenic (PS3+PM2+PM5+PP1+PP4) |
| 2: 44052029_A/T | Premature stop: Y301X | 4 | Pathogenic (PVS1+PS3) |
| 2: 44053544_G/A | Premature stop: Q251X | 7 | Pathogenic (PVS1+PS3) |
| 2: 44065689_A/C | Missense: S44A | 2 | Pathogenic (PS3+PM2+PM5+PP1+PP4) |
| ***ABCG8*** | | | |
| 2:44100970_TC/AA | Missense: I419K | 4 | Pathogenic (PS3+PM2+PM5+PP1+PP4) |
| 2:44100970_T/A | Missense: I419N | 9 | Pathogenic (PS3+PM2+PM5+PP3+PP4) |
| 2:44100999_A/G | Missense: M429V | 4 | Likely pathogenic (PS3+PM5+PP1+PP4) |

**Supplemental Table 2. Rare loss-of-function variants identified in *ABCG5*.**

| Variant | Consequence | MIGen, n | UK Biobank, n | TSCA, n |
|---|---|---|---|---|
| 2:44040325_G/C | Premature stop: S629X | 0 | 10 | 0 |
| 2:44040325_G/T | Premature stop: E629X | 0 | 4 | 0 |
| 2:44041643_C/A | Premature stop: E579X | 1 | 0 | 1 |
| 2:44041730_T/C | Splice Acceptor | 0 | 1 | 0 |
| 2:44041730_T/A | Splice Acceptor | 2 | 0 | 0 |
| 2:44047174_T/TG | Frameshift: H510fsTer | 0 | 0 | 1 |
| 2:44047211_G/A | Premature stop: R498X | 1 | 2 | 0 |
| 2:44047240_C/T | Splice Acceptor | 0 | 1 | 0 |
| 2:44047241_T/C | Splice Acceptor | 1 | 0 | 0 |
| 2:44049953_GA/G | Frameshift: F482fsTer | 1 | 0 | 0 |
| 2:44050063_G/A | Premature stop: R446X | 14 | 6 | 0 |
| 2:44050075_C/A | Splice Acceptor | 0 | 2 | 0 |
| 2:44050076_T/C | Splice Acceptor | 0 | 1 | 0 |
| 2:44051036_GCACGTGGG CACTTACA/G | Splice Donor | 4 | 0 | 0 |
| 2:44051051_C/A | Splice Donor | 1 | 0 | 0 |
| 2:44051357_C/G | Splice Donor | 1 | 0 | 0 |
| 2:44051431_T/A | Premature stop: K349X | 1 | 0 | 0 |
| 2:44052027_C/T | Splice Donor | 0 | 1 | 0 |
| 2:44052071_G/T | Premature stop: C287X | 2 | 0 | 0 |
| 2: 44052096_TC/T | Frameshift: Q279fsTer | 0 | 0 | 1 |
| 2:44052158_C/T | Splice Acceptor | 1 | 9 | 0 |
| 2:44053520_C/T | Splice Donor | 0 | 1 | 0 |
| 2:44053544_G/A | Stop Gained: Q251X | 0 | 1 | 0 |
| 2:44053568_G/A | Stop Gained: R243X | 3 | 0 | 0 |
| 2:44055121_C/G | Splice Donor | 1 | 0 | 0 |
| 2:44055121_C/T | Splice Donor | 1 | 0 | 0 |
| 2:44055180_GC/G | Frameshift: G192fsTer | 1 | 0 | 0 |
| 2:44055209_G/A | Stop Gained: R183Ter | 3 | 2 | 0 |
| 2:44058973_C/A | Stop Gained: E146Ter | 1 | 3 | 0 |
| 2:44059095_G/T | Stop Gained: Y131Ter | 1 | 0 | 0 |
| 2:44059154_C/A | Stop Gained: E131Ter | 0 | 1 | 0 |
| 2:44059166_TC/T | Frameshift: G107Ter | 4 | 0 | 0 |
| 2:44059223_C/T | Splice Acceptor | 0 | 1 | 0 |
| 2:44064975_G/C | Stop Gained: S88Ter | 1 | 2 | 0 |

| | | | | |
|---|---|---|---|---|
| 2:44065009_C/A | Stop Gained: E77Ter | 1 | 0 | 0 |
| 2:44065013_G/C | Stop Gained: Y75Ter | 0 | 2 | 0 |
| 2:44065051_G/A | Stop Gained: Q63Ter | 1 | 1 | 0 |
| 2:44065755_G/A | Stop Gained: Y45Ter | 5 | 0 | 0 |
| 2:44065684_G/C | Stop Gained: Q22Ter | 0 | 2 | 0 |
| Total | | 53 | 53 | 3 |

**Supplemental Table 3. Association of *ABCG8* loss-of-function variants with blood lipids and coronary artery disease risk among three WES cohorts.**

| Trait | LoF Carriers (n) | Total N | Effect (95% CI), P value |
|---|---|---|---|
| Total Cholesterol | 154 | 93049 | Beta 0.07 (-0.12 to 0.26), P = 0.49 |
| LDL Cholesterol | 154 | 93049 | Beta 0.08 (-0.06 to 0.23), P = 0.25 |
| HDL Cholesterol | 154 | 93049 | Beta -0.10 (-0.23 to 0.04), P = 0.16 |
| Triglycerides | 154 | 93049 | Beta 0.03 (-0.07 to 0.14), P = 0.58 |
| Coronary Artery Disease | 146 | 82915 | OR 0.79 (0.47 to 1.35), P = 0.39 |

Estimates include exome sequencing data from the Myocardial Infarction Genetics consortium and UK Biobank.

CI, 95% confidence interval; HDL, high-density-lipoprotein; LDL, low-density lipoprotein；LoF, loss-of-function; OR, odds ratio; SD, standard deviation; WES, whole-exome-sequencing.

**Supplemental Figure 1. Pedigrees of sitosterolemia families.**
Square and circle indicate male and female, respectively. Black or lattice pattern full half indicates heterozygote subjects; black, homozygote subjects; gray shading, genetically unknown subjects; and white, genetically unaffected subjects. Total cholesterol (mg/dL), triglycerides (mg/dL), high-density lipoprotein cholesterol (mg/dL), low-density lipoprotein cholesterol (mg/dL), and sitosterol (μg/mL) levels are displayed below each individual identifier. Patients who had a history of coronary artery disease are indicated with an asterisk "*".

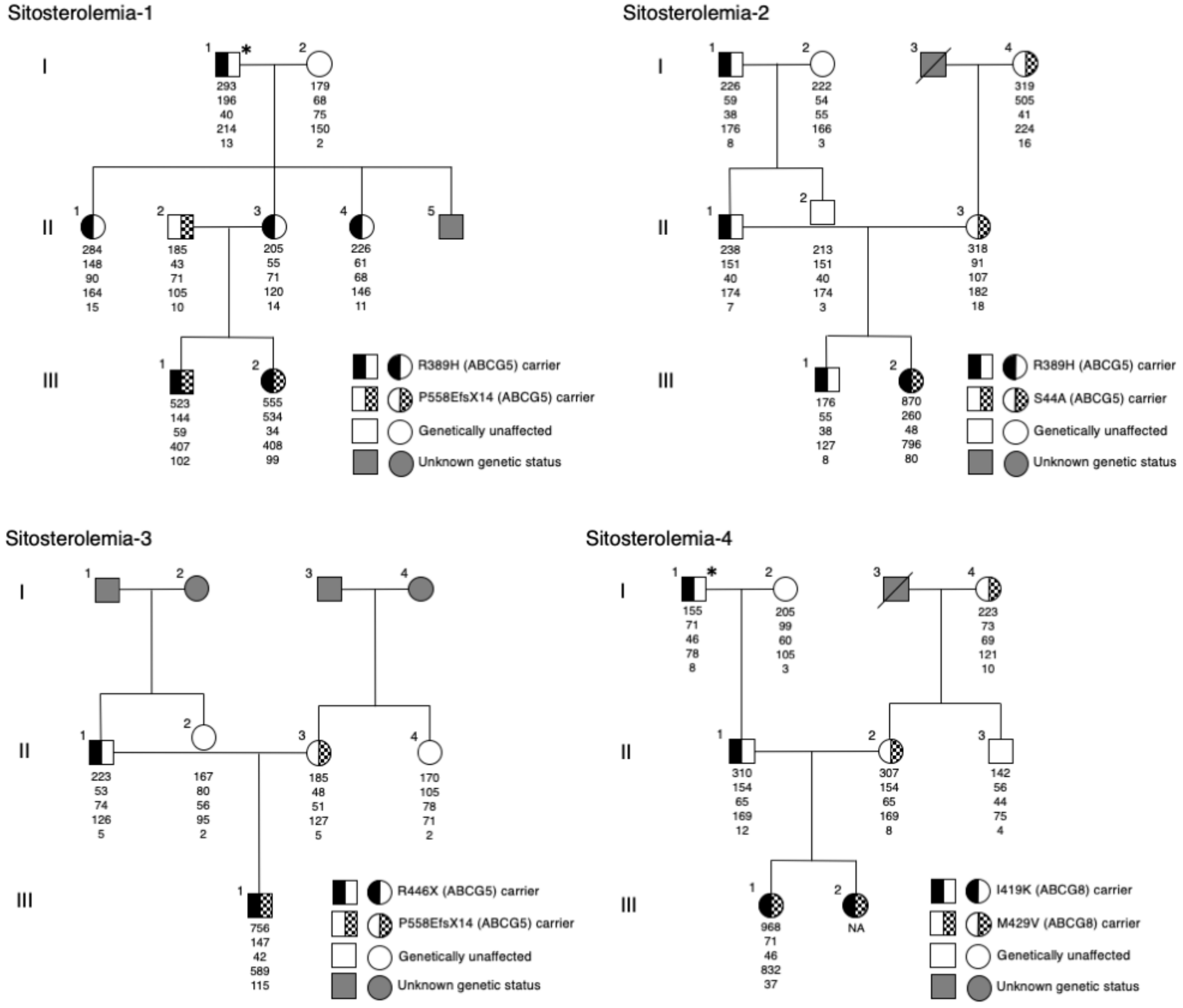

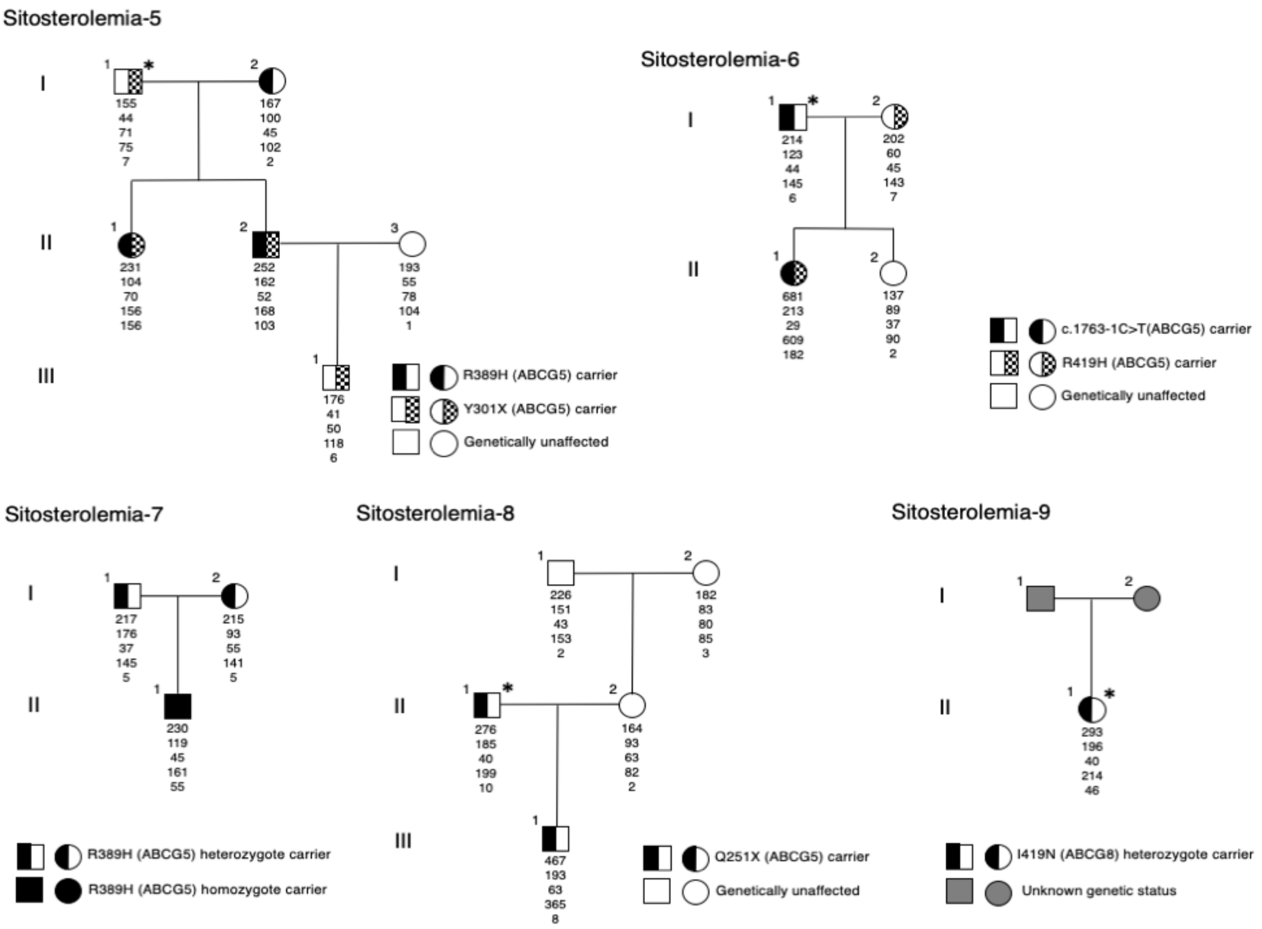

Sitosterolemia-5
I
II
III
155 44 71 75 7
167 100 45 102 2
231 104 70 156 156
252 162 52 168 103
193 55 78 104 1
176 41 50 118 6
R389H (ABCG5) carrier
Y301X (ABCG5) carrier
Genetically unaffected
Sitosterolemia-6
214 123 44 145 6
202 60 45 143 7
681 213 29 609 182
137 89 37 90 2
c.1763-1C>T(ABCG5) carrier
R419H (ABCG5) carrier
Genetically unaffected
Sitosterolemia-7
217 176 37 145 5
215 93 55 141 5
230 119 45 161 55
R389H (ABCG5) heterozygote carrier
R389H (ABCG5) homozygote carrier
Sitosterolemia-8
226 151 43 153 2
182 83 80 85 3
276 185 40 199 10
164 93 63 82 2
467 193 63 365 8
Q251X (ABCG5) carrier
Genetically unaffected
Sitosterolemia-9
293 196 40 214 46
I419N (ABCG8) heterozygote carrier
Unknown genetic status
